# Cross-Sampling Rate Transfer Learning for Enhanced Raw EEG Deep Learning Classifier Performance in Major Depressive Disorder Diagnosis

**DOI:** 10.1101/2023.11.13.566915

**Authors:** Charles A. Ellis, Robyn L. Miller, Vince D. Calhoun

## Abstract

Transfer learning offers a route for developing robust deep learning models on small raw electroencephalography (EEG) datasets. Nevertheless, the utility of applying representations learned from large datasets with a lower sampling rate to smaller datasets with higher sampling rates remains relatively unexplored. In this study, we transfer representations learned by a convolutional neural network on a large, publicly available sleep dataset with a 100 Hertz sampling rate to a major depressive disorder (MDD) diagnosis task at a sampling rate of 200 Hertz. Importantly, we find that the early convolutional layers contain representations that are generalizable across tasks. Moreover, our approach significantly increases mean model accuracy from 82.33% to 86.99%, increases the model’s use of lower frequencies, (θ-band), and increases its robustness to channel loss. We expect this analysis to provide useful guidance and enable more widespread use of transfer learning in EEG deep learning studies.

## 1. INTRODUCTION

Deep learning methods are increasingly being applied to raw resting-state electroencephalography (EEG) data. However, the development of reliable models can be challenging given the small size of many EEG datasets. Other deep learning application domains [1] often use transfer learning to boost model performance, and a handful of transfer learning approaches have been developed [2]–[4] for raw EEG. Moreover, some studies have suggested that the models they develop might be broadly applied to other datasets and conditions [3], [5]. One potential obstacle that could limit the widespread utility of these transfer learning approaches is the transfer of representations learned on data with a low sampling rate to models trained on data with higher sampling rates. In this study, we use an approach first developed in [5] and examine the utility of cross-sampling rate transfer learning. Importantly, we significantly boost the performance of a model trained for major depressive disorder (MDD) diagnosis by pretraining the model with a 100Hz sleep stage classification task.

Relative to machine learning and deep learning models trained on extracted features, deep learning models have the potential to identify important features that might otherwise be obscured [2]. As such, they are increasingly being applied to EEG tasks like MDD [5]–[7] and schizophrenia [8] diagnosis and emotion recognition [2]. Nevertheless, training robust deep learning models on raw EEG data can be difficult. Collecting large EEG datasets is often expensive and time-consuming [9], and deep learning models risk overfitting on small clinical EEG data [9]. One viable solution to this problem is the use of transfer learning.

Transfer learning has been widely applied within other deep learning domains like image classification [1], and a number of EEG studies have begun to develop transfer learning approaches for models trained on extracted features [10]–[12] or raw task data [13], [14]. Nevertheless, the application of transfer learning methods to deep learning models trained on raw, non-task EEG data remains relatively underdeveloped. Several studies have developed transfer learning approaches for raw resting-state EEG [2]–[5], and these studies suggest that pretrained EEG models like those in image classification [1] might one day see widespread use. While such an outcome could be undeniably beneficial for the field, there remain obstacles that could hamper the utility of pretrained standard models. One such obstacle is the effect of sampling rate upon the utility of transfer learning approaches. Many large, publicly available sleep datasets [15] that might be used for pretraining models have lower sampling rates than may be ideal for more complex tasks like disorder diagnosis [5]–[8]. To state that dilemma in question form, “Can the representations learned by a model trained upon a task with a low sampling rate enhance performance in another task with a higher sampling rate?”

In this study, we conduct a preliminary investigation into that question. Specifically, we (1) train a deep learning model on multichannel EEG data with a 200Hz sampling rate for MDD diagnosis using automated machine learning (AutoML), (2) train that architecture on a large single channel sleep stage dataset with a 100Hz sampling rate, (3) transfer the learned sleep representations to reinitialize the MDD model in a second round of training, and (4) apply spectral and spatial explainability approaches to examine the effect of the transfer learning approach upon the canonical frequency bands and channels prioritized by the model. We found that even with a cross-sampling rate transfer of features our approach increases model performance in a statistically significant manner.

## 2. METHODS

We describe our datasets, preprocessing, model development, transfer learning approach, and explainability analyses.

### 2.1 Datasets and Preprocessing

We used an MDD dataset to evaluate the utility of cross-sampling rate transfer learning and a sleep dataset to pretrain our models.

#### 2.1.1. MDD Dataset

We used an EEG dataset [16] of 5-to 10-minute resting-state recordings of 28 HCs and 30 MDDs with eyes closed that has been used in multiple deep learning studies [5]–[7], [17]. A sampling rate of 256 Hz and standard 10-20 format with 64 electrodes were used during recording. Like previous studies in MDD and other neuropsychiatric disorders [5], [6][18], we used 19 channels: Fp1, Fp2, F7, F3, Fz, F4, F8, T3, C3, Cz, C4, T4, T5, P3, Pz, P4, T6, O1, and O2. We downsampled the data to 200 Hz and separately channel-wise z-scored each recording. As a preliminary data augmentation approach to increase the number of available samples for training, we segmented the data into 25-second samples using a sliding window approach with a step size of 2.5 seconds. Our final MDD dataset had 2,942 MDDs and 2,950 HCs.

#### 2.1.1. Sleep Stage Dataset

To pretrain our models, we used the Sleep Cassette portion of the Sleep-EDF Expanded dataset [15] on PhysioNet [19]. The dataset had 153 20-hour recordings from 78 participants. It was recorded at a sampling rate of 100 Hz, and we used the Fpz-Cz electrode in our analysis. The data was assigned to Awake (AW), Rapid Eye Movement (REM, R), Non-REM 1 (N1), N2, N3, and N4 classes in 30-second segments by an expert. To reduce class imbalances, we removed Awake samples at the start and end of the recordings. Additionally, based on clinical guidelines [20], we combined N3 and N4 into a single N3 class. Lastly, we z-scored each recording separately. Our final dataset had 85,678, 21,497, 69,121, 13,039, and 25,832 AW, N1, N2, N3, and R samples, respectively.

### 2.2 Model Development Approach

We adapted an architecture (Figure 1) that was first presented in [21] and has been used for MDD classification [5]–[7]. However, unlike previous studies, we used the architecture as a starting point and then further optimized it using an AutoML approach in Keras-Tuner [22]. We trained a baseline MDD model – Model A – with Keras-Tuner and used the optimal architecture to train 8 more models. Model S was a sleep stage classification model. Models B through H were trained via transfer learning from Model S for MDD diagnosis. Unless otherwise specified, all MDD models used the same cross-validation and training approach. MDD model performance was quantified with accuracy (ACC), sensitivity (SENS), and specificity (SPEC), and Model S performance was quantified with the F1 score, SENS, and precision (PREC).

**Figure 1.**
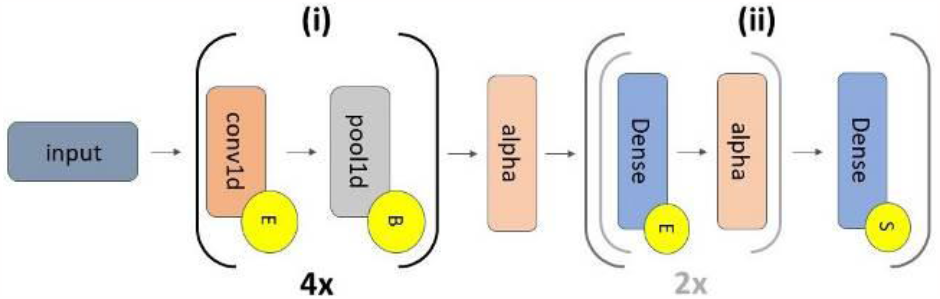
Model Architecture. All models have 2 sections separated by an alpha dropout layer (alpha) with a rate of 0.4: (i) a feature extraction portion, which repeats 4 times, and (ii) a classification portion. The grey subsection within (ii) repeats twice. The 4 convolutional (conv1d) layers have 10, 20, 5, and 20 filters, respectively with a kernel size of 20. They are followed by max pooling layers with a pool size of 2 and stride of 2. Section (ii) has 3 dense layers (16, 48, and 2 nodes) with interleaved alpha layers with dropout rates of 0.5. Yellow circles with an “E”, “B”, or “S” correspond to ELU activations, batch normalization, and softmax activations. All conv1d and dense layers had max norm kernel constraints with a max value of 1. In Model S, a channel dropout layer was located between “Input” and Section (i).

#### 2.1.1 Model A

We used the Hyperband algorithm in Keras-Tuner with 10 initial epochs per trial and a maximum of 40 training epochs. The Hyperband algorithm sought to optimize the average model validation ACC across folds. We used 25-fold subject-wise, stratified, group shuffle split cross-validation to have a sufficiently large number of folds to obtain statistically significant results and to prevent the mixing of samples from the same participant across training, validation, and test sets. Approximately, 70%, 20%, and 10% of the data was assigned to the training, validation, and test sets, respectively. To enhance baseline performance, we used data augmentation to triple the size of each training set. Our approach involved generating two duplicates of the training data and adding Gaussian noise with a mean of 0 and a standard deviation of 0.7 to one duplicate and a standard deviation of 0.5 to the other duplicate. We used Adam optimization (final learning rate = 0.001), a batch size of 128, and early stopping if model validation ACC did not increase following 5 subsequent training epochs. After hyperparameter tuning, we retrained with checkpoints to select models from the epochs with top validation ACC.

#### 2.1.2 Model S

When training Model S, we used the approach presented in [5]. Namely, we generated 18 duplicates of each 1-channel sleep sample, added Gaussian noise (μ=0, σ=0.7) to each duplicate, and combined the duplicates such that they had 19 channels like the MDD data. We then added a channel dropout layer [8] (rate=25%) to the start of the model to force it to learn from all input channels. We trained the model with 10-fold subject-wise, group shuffle split cross-validation with an 80-10-10 training, validation, test split, a maximum of 200 epochs, early stopping after 20 consecutive epochs without improved validation ACC, and an Adam optimizer (learning rate = 0.001) that decreased by an order of magnitude after 5 consecutive epochs without improved validation ACC.

#### 2.1.3 Models B, E, and G

We used the Model S weights to reinitialize the MDD model architecture. Upon initializing the architecture with the Model S weights, we froze some of the convolutional (conv) layers. In Model B, we froze all conv layers, and in models E and G, we froze the first conv layer and the first two conv layers, respectively. We used a virtually identical training and evaluation approach to that of Model A, with one modification. We repeated all 25 folds 10 times (i.e., one initialization per Model S fold). This created 250 models (i.e., 10 sleep folds x 25 MDD folds) per MDD model.

#### 2.1.4 Models C, F, and H

Initializing with the Model B, E, and G weights, we unfroze all conv layers and continued training with an identical crossvalidation approach to obtain Models C, F, and H, respectively. 2.1.4. Model D

As in Models B, E, and G, for Model D, we initialized with Model S weights. However, we did not freeze any layers during training. Model D had 250 models (i.e., 10 sleep folds x 25 MDD folds).

### 2.3 Statistical Comparison of Model Performance

To determine whether transfer learning significantly improved model performance, we generated 10 Model A performance duplicates (i.e., 1 duplicate per sleep fold) and performed familywise, paired, two-tailed t-tests between the ACC, SENS, and SPEC of each MDD model. We then applied false discovery rate (FDR) correction separately to the p-values of each metric.

### 2.4 Spectral Explainability Analysis

Given the cross-frequency nature of our transfer learning approach, we were curious how the importance of canonical frequency bands may have changed from the baseline to transfer learning-based models. As such, we applied a spectral explainability approach similar to that presented in [5]. Namely, we calculated the model ACC on the test samples in each fold, converted the test samples to the frequency domain with a Fourier transform, replaced the Fourier coefficients with values of zero within a frequency band of interest, converted back to the time domain, calculated the model ACC on the perturbed test samples, and computed the percent change (PCT_CHG) in model ACC. We analyzed the δ (0-4 Hz), θ (4-8 Hz), α (8-12 Hz), β (12-25 Hz), γ_1_ (25-45 Hz), γ_2_ (55-75 Hz), and γ_3_ (75-100 Hz) bands. We divided the γ-band into 3 subbands. γ_1_ and γ_2_ were divided due to 50Hz line noise, and γ_2_ and γ_3_ were divided to limit the effects of their perturbation to a ∼20 Hz band. We repeated this analysis for all folds of all MDD models.

### 2.5 Spatial Explainability Analysis

To examine how transfer learning affected individual channel importance, we applied a spatial explainability approach identical to that presented in [5]. Specifically, we ablated individual channels in the test data (i.e., replaced them with zeros) and calculated the PCT_CHG in ACC after ablation.

## 3. RESULTS AND DISCUSSION

In this section, we describe and discuss our model performance and explainability analysis results.

### 3.1 Model Performance Analysis

Tables 1 and 2 show the Model S and MDD model performance results, respectively. Figure 2 shows the results of our statistical analysis of model performance.

**Table 1.** Model S Performance.

|  | F1 | SENS | PREC |
| --- | --- | --- | --- |
| **AW** | 91.81 ± 01.92 | 93.97 ± 02.80 | 89.96 ± 04.36 |
| **N1** | 03.96 ± 01.31 | 02.15 ± 00.75 | 29.13 ± 06.40 |
| **N2** | 71.24 ± 05.52 | 62.88 ± 09.16 | 83.83 ± 05.47 |
| **N3** | 58.96 ± 12.60 | 91.29 ± 06.15 | 44.73 ± 13.49 |
| **R** | 54.99 ± 05.96 | 75.68 ± 09.63 | 43.79 ± 06.63 |

**Table 2.** MDD Model Performance.

| Model | ACC | SENS | SPEC |
| --- | --- | --- | --- |
| **A** | 82.33 ± 16.47 | 85.89 ± 24.22 | 79.49 ± 23.81 |
| **B** | 80.32 ± 13.95 | 80.40 ± 24.41 | 80.60 ± 21.17 |
| **C** | 80.41 ± 14.06 | 79.41 ± 24.80 | 81.77 ± 20.55 |
| **D** | 80.15 ± 14.19 | 80.23 ± 24.44 | 80.24 ± 21.32 |
| **E** | 85.17 ± 12.13 | 87.34 ± 17.70 | 81.62 ± 24.24 |
| **F** | **86.99 ± 12.57** | 87.55 ± 20.13 | **85.27 ± 24.31** |
| **G** | 83.40 ± 13.47 | **88.97 ± 17.62** | 76.14 ± 27.23 |
| **H** | 86.50 ± 12.39 | 87.55 ± 20.02 | 84.79 ± 24.12 |

**Figure 2.**
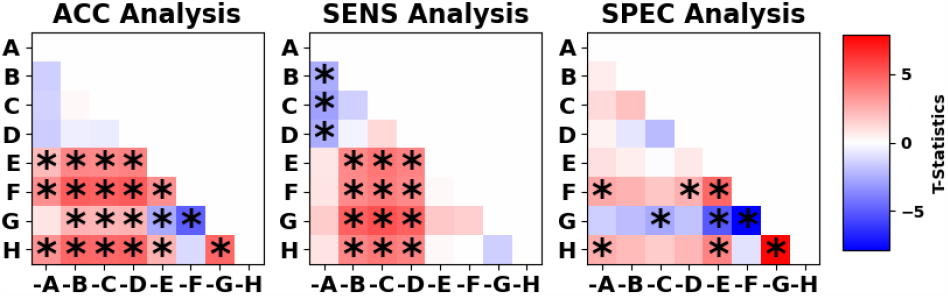
Model Performance Analysis. The panels show differences in ACC, SENS, and SPEC from left to right, respectively. The y-axis shows the MDD models, and the x-axis shows the models subtracted from the models on the y-axis. T-statistics for each t-test are shown on the heatmaps, and asterisks show those with significance after correction (p < 0.05). All panels share the same color bar.

#### 3.1.1 Model S

Model S performance was, with the exception of N1, well above chance-level. While Model S obtained lower performance than some sleep classification studies [23], high Model S performance was not the goal of our study. The goal of Model S was to provide a more domain-informed initialization for our MDD models.

#### 3.1.2 MDD Models

The performance of Model A, our baseline model, was competitive with other MDD studies on the same dataset [6], [7], and our use of automated hyperparameter tuning suggests that we likely obtained near peak performance for our dataset and architecture. Initializing with Model S weights and training with all conv layers unfrozen (Model D) or with all conv layers frozen (Model B) and subsequently unfrozen (Model C) decreased model performance. This hints at the effect of cross-sampling rate transfer learning, as these approaches yielded significant performance improvements in a previous EEG transfer learning study on data with the same sampling rates [5]. This insight, combined with how our models trained with frozen early conv layers (Models E and G) followed by unfrozen training (Models F and H) obtained significant improvements in ACC relative to Model A, suggests that the lower level feature representations learned in the early model layers were more useful in cross-sampling rate transfer learning than the higher-level features extracted in later layers. It further demonstrates that our approach has the potential for significant utility in cross-sampling rate EEG transfer learning tasks.

### 3.2 Spectral Explainability Analysis

Figure 3 shows our spectral explainability results. The results are generally similar across models; δ and β had high importance. Interestingly, in Model A, θ was of low importance. However, Models E through H seemed to use θ effectively, which could explain their increase in performance relative to Models A through D. This could be an effect of cross-sampling rate transfer learning. Given the 4-7 Hz frequency range of θ at 200 Hz, the identified waveforms would have likely appeared at 8-14 Hz (α) in the 100 Hz sleep data, which would roughly correspond to the 11-16 Hz sleep spindles of N2 [20]. This is also interesting given that θ has been important for MDD classification [6], [7], [24] in some studies but not others [5]. Across MDD models, α and γ had relatively minor, but non-trivial, effects. Perturbation of γ_1_ perturbation had a minor effect in Models B through D and a smaller effect on Models F through H. Among the pretrained models, α perturbation most affected Models E and G, which conforms to the identified importance of α in previous studies [25].

**Figure 3.**
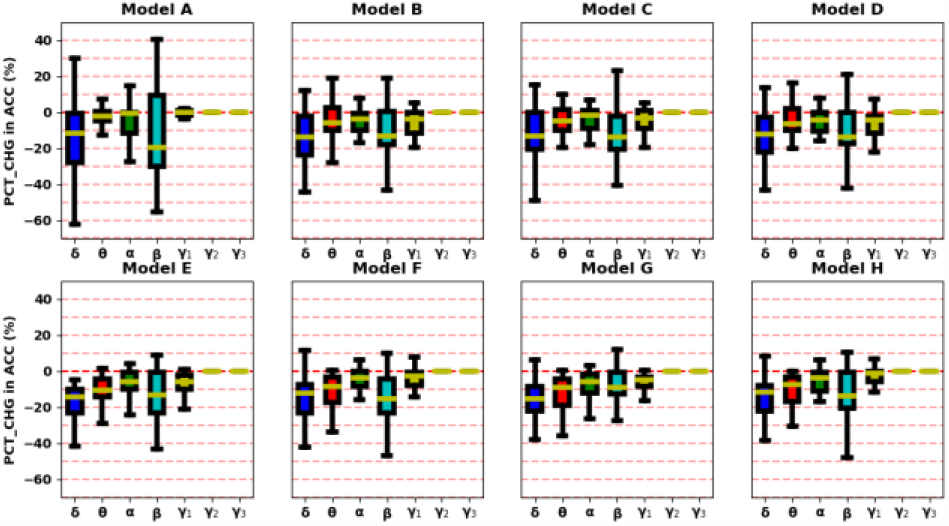
Spectral Importance. The top and bottom panels show Models A through D and E through H, respectively. To mediate the effects of variance resulting from having 25 models for Model A and 250 Models for Models B through H, we averaged the mean PCT_CHG in ACC for each set of 10 sleep folds. The y-axes of the leftmost panels coincide with those of the other panels. The x-axes indicate frequency bands.

### 3.3 Spatial Explainability Analysis

Figure 4 shows our spatial explainability results. Interestingly, our models seemed to be separated into three groups based on their spatial explanations: (1) Model A, (2) Models B through D, and (3) Models E through H. Model A most prioritized F7 and Cz. In alignment with previous studies [5], Models B through H prioritized Fz, Pz, and O1, with some importance on C3, Cz, and C4. Lastly, channel perturbation in Models E through H did not seem to have any major effect upon model performance. This likely indicates that the channel-wise representations learned on the sleep data in the early model layers were very robust (possibly due to our use of channel dropout) but that the remainder of the model needed to adjust to effectively use those representations.

**Figure 4.**
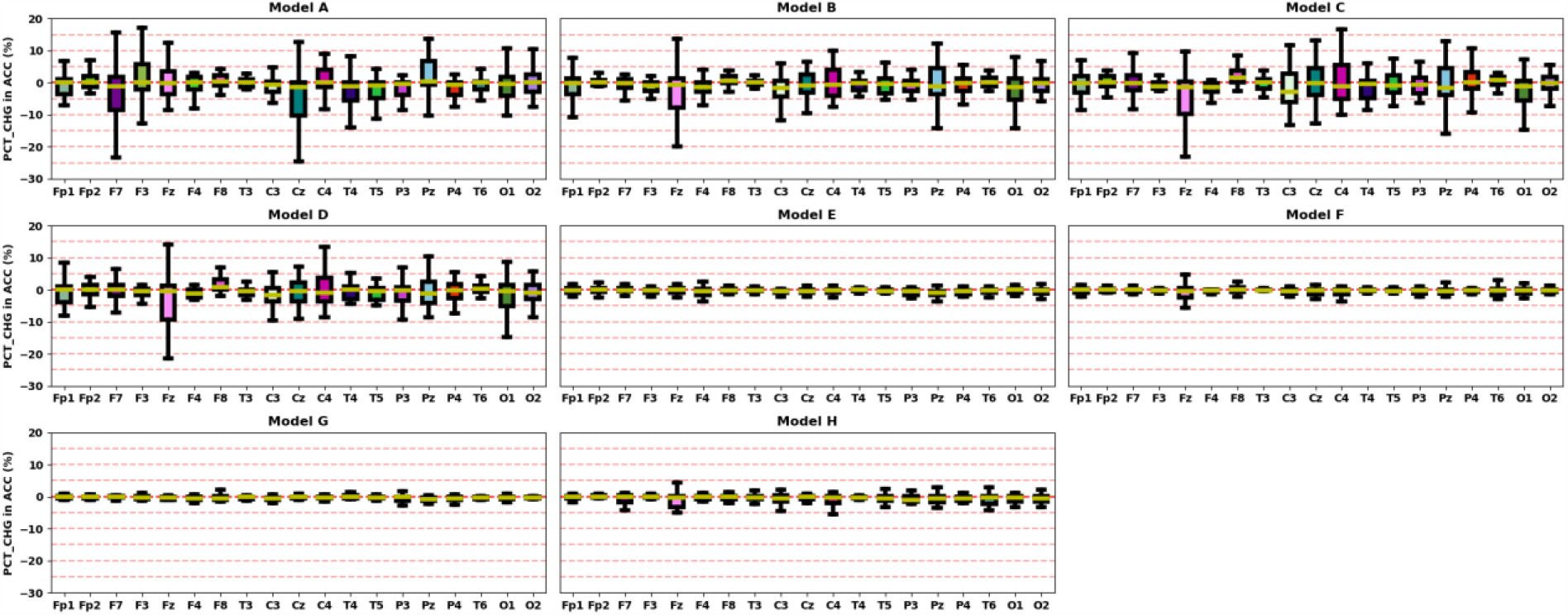
Spatial Explainability Results. Each panel shows the results for a different model. To mediate the effects of variance resulting from having 25 models for Model A and 250 Models for Models B through H, we averaged the mean PCT_CHG in ACC for each set of 10 sleep folds. The y-axes of the leftmost panels coincide with those of the other panels. The x-axes indicate channels.

### 3.4 Limitations and Future Work

Due to computational constraints with Keras-Tuner, we did not optimize Model S hyperparameters (e.g., learning rate or number of dense layer nodes). However, further hyperparameter tuning on the sleep data could yield better representations. To minimize the number of confounders in our analysis, we used the same learning rate and dense layer architecture in MDD Models B through H that we used in Model A. Further hyperparameter tuning with Model S initializations could potentially enhance model performance. Future studies might also examine the effect of transfer learning on the important of waveforms [23]. Additionally, applying the approach to other EEG datasets and classification tasks (e.g., schizophrenia diagnosis [18]) would help validate its utility.

## 4 CONCLUSION

Developing robust deep learning models trained on raw EEG data can be challenging due to limited data availability. Moreover, transfer learning approaches could enable the development of more robust models on smaller datasets, but if they are to be implemented in a widespread manner, the utility of models pretrained on data with lower sampling rates for transfer to tasks with data of higher sampling rates needs to be further investigated. In this study, we show that representations encoded in early convolutional layers in a sleep stage classification task can significantly boost performance in a higher sampling rate MDD diagnosis task. Furthermore, we show that the approach increases the importance of lower frequency bands to model performance and increases the robustness of the model to channel loss. We hope that this study will inform future EEG transfer learning developments and help enable their widespread implementation.

## 5. ACKNOWLEDGMENTS

This study was funded by NIH R01MH123610, NIH R01MH118695, and NSF 2112455.

## 6. COMPLIANCE WITH ETHICAL STANDARDS

Given that both MDD and sleep datasets were publicly available, it was unnecessary to obtain IRB approval for the study.

## Notes

https://www.physionet.org/content/sleep-edfx/1.0.0/

https://figshare.com/articles/dataset/EEG_Data_New/4244171

### Competing Interest Statement

The authors have declared no competing interest.

## REFERENCES

[1] K. Simonyan and A. Zisserman, “Very Deep Convolutional Networks for Large-Scale Image Recognition,” in International Conference on Learning Representations (ICLR), 2015, pp. 1–14.

[2] Y. Cimtay and E. Ekmekcioglu, “Investigating the use of pretrained convolutional neural network on cross-subject and cross-dataset eeg emotion recognition,” Sensors (Switzerland), vol. 20, no. 7, pp. 1–20, 2020, doi: 10.3390/s20072034.

[3] P. Nejedly et al., “Intracerebral EEG Artifact Identification Using Convolutional Neural Networks,” Neuroinformatics, vol. 17, no. 2, pp. 225–234, 2019, doi: 10.1007/s12021-018-9397-6.

[4] H. Phan, O. Y. Chén, P. Koch, A. Mertins, and M. De Vos, “Deep transfer learning for single-channel automatic sleep staging with channel mismatch,” Eur. Signal Process. Conf., vol. 2019-Septe, 2019, doi: 10.23919/EUSIPCO.2019.8902977.

[5] C. A. Ellis, R. L. Miller, and V. D. Calhoun, “Improving Multichannel Raw Electroencephalography-based Diagnosis of Major Depressive Disorder via Transfer Learning with Single Channel Sleep Stage Data,” 2023.

[6] C. A. Ellis, A. Sattiraju, R. L. Miller, and V. D. Calhoun, “Novel Approach Explains Spatio-Spectral Interactions in Raw Electroencephalogram Deep Learning Classifiers,” 2023.

[7] C. A. Ellis, A. Sattiraju, R. L. Miller, and V. D. Calhoun, “A Framework for Systematically Evaluating the Representations Learned by A Deep Learning Classifier from Raw Multi-Channel Electroencephalogram Data,” bioRxiv, 2023.

[8] A. Sattiraju, C. A. Ellis, R. L. Miller, and V. D. Calhoun, “An Explainable and Robust Deep Learning Approach for Automated Electroencephalography-based Schizophrenia Diagnosis,” 2023.

[9] Z. Wan, R. Yang, M. Huang, N. Zeng, and X. Liu, “A review on transfer learning in EEG signal analysis,” Neurocomputing, vol. 421, pp. 1–14, 2021, doi: 10.1016/j.neucom.2020.09.017.

[10] M. T. Sadiq, M. Z. Aziz, A. Almogren, A. Yousaf, S. Siuly, and A. U. Rehman, “Exploiting pretrained CNN models for the development of an EEG-based robust BCI framework,” Comput. Biol. Med., vol. 143, no. August 2021, p. 105242, 2022, doi: 10.1016/j.compbiomed.2022.105242.

[11] P. Jadhav, G. Rajguru, D. Datta, and S. Mukhopadhyay, “Automatic sleep stage classification using time–frequency images of CWT and transfer learning using convolution neural network,” Biocybern. Biomed. Eng., vol. 40, no. 1, pp. 494–504, 2020, doi: 10.1016/j.bbe.2020.01.010.

[12] A. Vilamala, K. H. Madsen, and L. K. Hansen, “Deep convolutional neural networks for interpretable analysis of EEG sleep stage scoring,” IEEE Int. Work. Mach. Learn. Signal Process. MLSP, vol. 2017-Septe, no. 659860, pp. 1–6, 2017, doi: 10.1109/MLSP.2017.8168133.

[13] M. Azab, H. Ahmadi, L. Mihaylova, and M. Arvaneh, “Dynamic time warping-based transfer learning for improving common spatial patterns in brain-computer interface,” J. Neural Eng., vol. 17, no. 1, 2020, doi: 10.1088/1741-2552/ab64a0.

[14] Ditthapron, N. Banluesombatkul, S. Ketrat, E. Chuangsuwanich, and T. Wilaiprasitporn, “Universal Joint Feature Extraction for P300 EEG Classification Using Multi-Task Autoencoder,” IEEE Access, vol. 7, pp. 68415–68428, 2019, doi: 10.1109/ACCESS.2019.2919143.

[15] “PhysioNet: The Sleep-EDF database [Expanded].”

[16] W. Mumtaz, L. Xia, M. A. M. Yasin, S. S. A. Ali, and A. S. Malik, “A wavelet-based technique to predict treatment outcome for Major Depressive Disorder,” PLoS One, vol. 12, no. 2, pp. 1–30, 2017, doi: 10.1371/journal.pone.0171409.

[17] H. W. Loh, C. P. Ooi, E. Aydemir, T. Tuncer, S. Dogan, and U. R. Acharya, “Decision support system for major depression detection using spectrogram and convolution neural network with EEG signals,” Expert Syst., vol. 39, no. 3, pp. 1–15, 2022, doi: 10.1111/exsy.12773.

[18] A. Ellis, A. Sattiraju, R. Miller, and V. Calhoun, “Examining Effects of Schizophrenia on EEG with Explainable Deep Learning Models,” in 2022 IEEE 22nd International Conference on Bioinformatics and Bioengineering (BIBE), 2022, pp. 301–304. doi: 10.1109/BIBE55377.2022.00068.

[19] G. AL et al., “PhysioBank, PhysioToolkit, and PhysioNet: Components of a New Research Resource for Complex Physiologic Signals,” Circulation, vol. 101, no. 23, pp. e215–e220, 2000, [Online]. Available: http://circ.ahajournals.org/content/101/23/e215.full

[20] Iber, S. Ancoli-Israel, A. L. Chesson, and S. F. Quan, “The AASM Manual for Scoring of Sleep and Associated Events: Rules, Terminology, and Technical Specifications.” 2007.

[21] S. L. Oh, J. Vicnesh, E. J. Ciaccio, R. Yuvaraj, and U. R. Acharya, “Deep convolutional neural network model for automated diagnosis of Schizophrenia using EEG signals,” Appl. Sci., vol. 9, no. 14, 2019, doi: 10.3390/app9142870.

[22] T. O’Malley, E. Bursztein, J. Long, F. Chollet, H. Jin, and L. Invernizzi, “KerasTuner,” 2019.

[23] A. Ellis, R. L. Miller, and V. D. Calhoun, “A Model Visualization-based Approach for Insight into Waveforms and Spectra Learned by CNNs,” in Proceedings of the Annual International Conference of the IEEE Engineering in Medicine and Biology Society, EMBS, 2021, pp. 1–4.

[24] R. A. Movahed, G. P. Jahromi, S. Shahyad, and G. H. Meftahi, “A major depressive disorder classification framework based on EEG signals using statistical, spectral, wavelet, functional connectivity, and nonlinear analysis,” J. Neurosci. Methods, vol. 358, no. November 2020, p. 109209, 2021, doi: 10.1016/j.jneumeth.2021.109209.

[25] A. A. Fingelkurts, A. A. Fingelkurts, H. Rytsälä, K. Suominen, E. Isometsä, and S. Kähkönen, “Composition of brain oscillations in ongoing EEG during major depression disorder,” Neurosci. Res., vol. 56, no. 2, pp. 133–144, 2006, doi: 10.1016/j.neures.2006.06.006.

